# Immunoinformatic Designing and Evaluation of a Broad-Spectrum Multiepitope Vaccine Against MDR *Acinetobacter baumannii, Klebsiella pneumoniae* and *Pseudomonas aeruginosa*

**DOI:** 10.1101/2025.05.27.656513

**Authors:** Neha Bhardwaj, Shiv Nandan Sah, Anshika Billa, Varsha Gupta, Neena Capalash, Prince Sharma

## Abstract

*Acinetobacter baumannii, Klebsiella pneumoniae* and *Pseudomonas aeruginosa* are among the multidrug-resistant (MDR) Gram-negative pathogens that pose a growing threat, necessitating novel preventive measures in addition to traditional antibiotics. By using advanced immunoinformatics methods, highly conserved and immunogenic epitopes for B-cells and T-cells were selected from major virulence-associated proteins (LptE, YiaD, MrkD, PhoE, OprF, and Zot), which exhibited high antigenicity (VaxiJen scores 0.74-2.75) and 98.87% global population coverage. The construct vaccine comprises 50S ribosomal protein L7/L12 adjuvant and PADRE sequence for immunogenicity enhancement, with structural validation indicating stability (96.3% residues in the total allowed Ramachandran regions). High-affinity interactions with TLR2/TLR4 (binding energies: -1009.6 to -1079.6 kcal/mol) were found through molecular docking, and immune simulations suggested strong humoral (IgM/IgG) and cellular (IFN-γ/IL-12) responses. Importantly, MEP vaccines can overcome major drawbacks of traditional vaccines by (1) offering cross-strain protection via conserved epitopes, (2) lowering the need for antibiotics through infection prevention, and (3) providing affordable options for healthcare systems affected by MDR infections. These findings demonstrate the MEP construct’s potential as a preventative measure against nosocomial infections, which may have implications for combating the global AMR epidemic. Further experimental validation is needed to verify its efficacy.

## Introduction

MDR bacteria represent one of the twenty-first century’s most important global health issues. The careless abuse of antibiotics has sped up antibiotic resistance worldwide, resulting in the emergence of “superbugs” that kill millions of people every year (Parmanik et al., 2022). MDR pathogens are a major public health concern that, if unchecked, as per WHO, it could result in up to 1 crore annual deaths by 2050, according to the World Health Organization (WHO). (Pulingam et al., 2022). ESKAPE pathogens, which include *Enterococcus faecium, Staphylococcus aureus, Klebsiella pneumoniae, Acinetobacter baumannii, Pseudomonas aeruginosa* and *Enterobacter spp*., are especially known to evade medications and cause nosocomial infections that can be fatal (Miller et al., 2024).

*A. baumannii, K. pneumoniae*, and *P. aeruginosa* use different resistance mechanisms, including integron-mediated gene transfer, biofilm formation, efflux pumps, outer membrane protein changes, and β-lactamase overexpression (Thomson et al., 2005). Carbapenem-resistant Enterobacterales (CRE) and Carbapenem-resistant

*A. baumannii* (CRAB) are highlighted in the WHO’s 2024 Bacterial Priority Pathogen List as “critical” threats, while *P. aeruginosa* (CRPA) was reclassified to “high” priority due to evolving resistance patterns (WHO, 2024). The widespread occurrence of these strains in medical environments highlights the pressing need for substitute treatment approaches (Chang et al., 2021).

Immunisation is a highly effective and cost-efficient strategy for infectious disease prevention (Sharma et al., 2016). Preventive immunization of high-risk groups, such as intensive care unit patients, may help reduce MDR nosocomial infections. Subunit vaccines (recombinant proteins, polysaccharide conjugates), whole-organism vaccines (live-attenuated or inactivated), and advanced platforms such as bacterial or viral vectors (Cagigi et al., 2023), mRNA/DNA vaccines (Pollard et al., 2021), and nanoparticle-based delivery systems are examples of current vaccine strategies.

Molecular docking analyses, immune response simulation, and the accurate identification of immunodominant epitopes have all been made possible by recent developments in computational immunology, which have completely changed the design of vaccines (Guarra et al., 2023). *In silico* tools (e.g., BepiPred, NetMHC, VaxiJen) allow for the design of multiepitope chimeric antigens with optimal immunogenicity, solubility, and safety profiles (Barman et al., 2022). Such vaccines can target conserved virulence factors across multiple pathogens, offering broad-spectrum protection (Douradinha et al., 2024). Clinically, MEP vaccines can reduce MDR infection rates, shorten hospital stays, and lower healthcare costs, particularly in high-risk settings like ICUs. Most importantly, by pre-emptively blocking infections, they alleviate the financial burden on healthcare systems caused by prolonged MDR treatments, offering a cost-effective, proactive solution to the global antibiotic resistance crisis (Ullah et al., 2024).

This research aims to create a new synthetic vaccine based on epitopes that will protect against *A. baumannii, K. pneumoniae*, and *P. aeruginosa* by identifying conserved, immunogenic epitopes using reverse vaccinology (Singh et al., 2016; Singh et al., 2017), designing a chimeric antigen with adjuvants to enhance immune responses, and validating the construct through *in silico* immunogenicity and safety analyses. In addition to their extensive population coverage, which resolves immune response differences across demographics, they overcome strain variability by adding conserved epitopes, ensuring cross-protection across various bacterial variants.

In this study, a total of six proteins, essential for their respective nosocomial infection-causing bacteria (*A. baumannii, K. pneumoniae*, and *P. aeruginosa*) that are reported as significant vaccine candidates, were selected, viz. LptE (Lipopolysaccharide transport E), a LPS transport scaffold vital for maintaining the integrity of the cell envelope, the average antigenic score demonstrated it to be a viable vaccination candidate targeting *A. baumannii* (Beiranvand *et. al*., 2021), YiaD (outer membrane porin), required for adhesion as well as for biofilm formation and is a surface antigen that is highly conserved and capable of eliciting a protective immune response against *A. baumannii* (Hagag *et. al*., 2022). MrkD (Type 3 Fimbrial adhesion protein), essential for biofilm formation of *K. pneumoniae* on biotic/abiotic surfaces (catheters, ventilators) (Ragupathi et. al., 2024), PhoE, a highly conserved outer membrane phosphoporin that regulates phosphate transport under nutrient limitation and is a potential candidate antigen for vaccine against *K. pneumoniae* infection (Hu *et. al*., 2022). OprF, a major outer membrane porin that maintains structural integrity, is critical for biofilm maturation and antibiotic tolerance and is also a standalone vaccine candidate for *P. aeruginosa* (Bahey *et. al*., 2020) and Zot (Zonula occludens toxin), a tight junction disruptor that helps in mucosal barrier penetration of host tissue, which makes it a “high value” vaccine candidate that can fight off all genetic variants of *P. aeruginosa* by triggering polarized immunological response (Benyamini *et. al*., 2024).

## Materials and Methods

From each of the bacteria—*A. baumannii, K. pneumoniae*, and *P. aeruginosa*—two vaccine candidates were chosen, respectively, to construct the MEP, viz. LptE (UniProt ID: A0A009PDL7), YiaD (UniProt ID: A0A5K6CUN9), MrkD (UniProt ID: P21648), PhoE (UniProt ID: P30704), OprF (UniProt ID: P37726), Zot (UniProt ID: B3G280), their sequence were obtained from UniProt (http://www.uniprot.org/) in FASTA format (Sah *et. al*., 2024).

### Conservation analysis

Conservation of the vaccine candidate proteins from *A. baumannii, K. pneumoniae*, and *P. aeruginosa* was analyzed with *K. pneumoniae* and *P. aeruginosa* and vice versa, and also in other strains of *A. baumannii, K. pneumoniae*, and *P. aeruginosa*, respectively, by BLAST, a tool from NCBI (http://blast.ncbi.nlm.nih.gov/Blast.cgi). A conservation analysis was done based on sequence alignment, showing QC and PI. BLASTn and BLASTp tools were used for nucleic acid and amino acid sequences, respectively (Touhidinia *et al*. 2021). On the basis of conservation analysis results of the selected proteins, LptE was selected as the most conserved among all the selected proteins.

### Topology prediction

The BOCTOPUS2 server (https://b2.topcons.net/) was used for predicting the topology of LptE. This web server can find the topology of transmembrane β-strands, inner loops, and outer loops in long sequences. Prediction of outer loops determines which amino acids are probably located outside the membrane (Hayat *et. al*., 2016). The sequence of 120 amino acids from the outer membrane loop region was selected and used as a backbone for the MEP construct (Ren *et. al*., 2019).

### Selection of suitable predicted B cell and T cell epitopes for designing the MEP

The IEDB server was utilized to predict B-cell and T-cell epitopes. Linear B-cell epitopes were identified using the Bepipred 2.0 tool (http://tools.iedb.org/bcell/) with a cutoff score of 0.5 (Sah et al., 2024). For MHC class I epitope prediction, the Artificial Neural Network (ANN) 4.0 algorithm (http://tools.iedb.org/mhci/) was employed, while MHC class II epitopes were predicted via the NN-align 2.3 (NetMHCII 2.3) method (http://tools.iedb.org/mhcii/) (Mubarak et. al., 2024).

### Epitope antigenicity, toxicity, and allergenicity prediction

Potential B-cell epitopes and T-cell epitopes from each protein were screened based on important parameters, including antigenicity, toxicity, and allergenicity. Antigenicity was assessed using the VaxiJen (v2.0) server (https://www.ddg-pharmfac.net/vaxijen/VaxiJen/VaxiJen.html), while allergenicity was evaluated with AllerTOP (https://www.ddg-pharmfac.net/allertop_test/). Toxicity predictions were performed using ToxinPred (https://webs.iiitd.edu.in/raghava/toxinpred/multi_submit.php). (Ayyagari, 2023).

### Population Coverage

For population coverage calculations, the specialized immunoinformatics tool available through the Immune Epitope Database platform (IEDB; http://tools.immuneepitope.org/tools/population/iedb_input) ensures effectiveness of the designed vaccine for the maximum proportion of the global human population for unique expected epitopes (Fleri et al. 2017). Each of the epitope and the matching binding HLA alleles were added, and different geographical areas were chosen in order to determine the vaccine’s population coverage.

### MEP vaccine design

The sequence of LptE was used as a backbone for the MEP construct where the shortlisted epitopes were inserted into the outer membrane loop region of LptE (Ren *et. al*., 2019). The selected immunogenic and conserved B-cell and T-cell (MHC-I and MHC-II) epitopes were connected by suitable linkers. Various linkers like Ala-Ala-Tyr (AAY) for MHC-I epitopes, GlyPro-Gly-Pro-Gly (GPGPG) for MHC-II epitopes, and Lys-Lys (KK) for the linear B-cell epitopes, respectively, were used. GlyPro-Gly-Pro-Gly (GPGPG) was utilised to connect the chosen MHC-I epitopes, Lys-Lys (KK) was incorporated to join the epitopes from B-cell, and Ala-Ala-Tyr (AAY) linkers joined MHC-II epitopes. Also, at the construct’s N-terminal, an EAAAK linker was used to connect a strong Pan DR (PADRE; CD4 activator) after an adjuvant (Abdus *et al*., 2020).

The sequential order of epitopes in different arrangements was identified through the process of epitope shuffling. Based on analytical investigations, the best configuration was selected (Sah *et. al*., 2024). Four different types of constructs by incorporating a distinct adjuvant into each vaccine sequence were created, each containing a distinct adjuvant: GABA protein, 50S ribosomal protein L7/L12, β-defensin, and Cholera Toxin B subunit (CTBS).

### Physiochemical characterisation

Molecular properties of vaccine construct, such as its molecular mass, amino acid composition, predicted pI, stability matrics, grand average of hydropathicity (GRAVY), and aliphatic index, were analyzed using ProtParam (https://web.expasy.org/protparam/).

VaxiJen (v2.0) was used to analyze each construct’s virulence, toxicity, antigenicity, and allergenicity. Allertop (https://www.ddg-pharmfac.net/allertop_test/) for allergenicity, VirulentPred (https://bioinfo.icgeb.res.in/virulent2/index.html) for virulence, and for antigenicity, VaxiJen (http://www.ddg-pharmfac.net/vaxijen/VaxiJen/VaxiJen.html) were used. The toxicity will be assessed using ToxinPred (https://webs.iiitd.edu.in/raghava/toxinpred/multi_submit.php) (Dimitrov *et. al*.,2013; Gupta *et. al*., 2013).

### Prediction of secondary structure

Protein secondary structural components were derived using PSIPRED (http://bioinf.cs.ucl.ac.uk/psipred/), which analyzes protein sequence segments using the Protein Data Bank (PDB) library to identify conserved substructures. Additionally, secondary structural components were validated using SOPMA (https://npsaprabi.ibcp.fr/cgibin/npsa_automat.pl?page=/NPSA/npsa_sopma.html) and GOR IV (https://npsa-prabi.ibcp.fr/cgi-bin/npsa_automat.pl?page=/NPSA/npsa_gor4.html) (Bibi *et al*., 2021).

### Prediction of tertiary structure

To forcast the tertiary structure I-TASSER tool (https://zhanggroup.org/I-TASSER/) was used. For visualizing the tertiary structure Discovery Studio Visualizer was utilized (Feig *et al*., 2017). In order to create a structure that was comparable to the experimental structure GalaxyRefine (http://galaxy.seoklab.org/cgi-bin/submit.cgi?type=REFINE) was used to modify the first developed tertiary structure model in a precise and superior way. By optimizing side chains and employing molecular dynamic simulations, the intended MEP 3D structural quality was enhanced, producing a more refined model (Hu *et al*. 2019). By creating Ramachandran plots for every model using PROCHECK in PDBsum (https://www.ebi.ac.uk/thorntonsrv/databases/pdbsum/Generate.html), the 3D structure of every construct was confirmed (Sah et al. 2024). These criteria were used to choose a final vaccine design for additional research.

### Prediction of conformational B cell epitope

Ellipro (http://tools.iedb.org/ellipro/) was used to anticipate the MEP construct’s conformational B-cell (Jaiswal *et al*., 2020)

### Docking studies

ClusPro v.2 online server (Khalid et al., 2022) was used for molecular docking. It is an automated web-based docking program that can employ a multi-stage process that includes rigid PIPER docking, filtering and clustering of docked conformations, and stabilizing using Monte Carlo simulations between the final MEP construct and host immune cell receptors like TLR2 (PDB ID: 2Z7X) and TLR4 (PDB ID: 3FXI) (Desta et al. 2020). Based on their binding energies, the results were grouped. Toll-like receptors from the NCBI Pubchem library and PDB database were downloaded, and they were docked with the vaccine construct using the Cluspro 2/Patchdock online docking tool (Sah *et al*., 2024)

### Codon optimization and In-silico cloning

The vaccine’s amino acid sequence was reverse-translated into its corresponding gene (cDNA) using the EMBOSS Transeq tool (https://www.ebi.ac.uk/jdispatcher/st/emboss_backtranseq). To improve the expression of heterologous proteins in the E. coli host strain (K12), the codons of the translated MEP gene were modified using the Java Codon Adaptation Tool (JCAT) (Grote et al. 2005). Codon Adaptation Index (CAI) values and GC content, avoiding bacterial ribosome-binding sites, restriction enzyme cleavage sites, and Rho-independent transcription termination, were the criteria for modifying the gene’s codon (Bibi et al. 2021). Using the Snapgene tool, the final vaccine construct with modified codons was cloned into the E. coli pET-28a (+) vector to guarantee the expression of the vaccine design.

### Immune Simulation Analysis

C-ImmSim (https://kraken.iac.rm.cnr.it/C-IMMSIM) was applied to evaluate the human body’s immune response. It produced models of humoral and cellular immune responses that were comparable to *in vivo* investigations (Rapin *et al*. 2012).

## Results

### Screening of immunodominant B-cell, T-cell (MHC-I and MHC-II) epitopes

LptE was selected as the backbone of MEP, and its topology was predicted using the BOCTOPUS server, which showed the outer loop region in its sequence (Fig. 2(a)). Based on this, the site for the insertion of B-cell and T-cell epitopes from YiaD, MrkD, PhoE, OprF and Zot was determined. One potential B-cell epitope was identified from each selected immunogenic protein. The IEDB server predicted the MHC-I and MHC-II epitopes for each of these proteins. The epitopes that satisfied the following criteria were screened for additional analysis: potent immunogenic characteristics, such as non-allergens with high antigenicity, low IC50 score (≥100), and high binding scores. Additionally, from each vaccine candidate, only the MHC-I and MHC-II epitopes that had the highest antigenicity score were kept (Table 1).

**Table 1:**
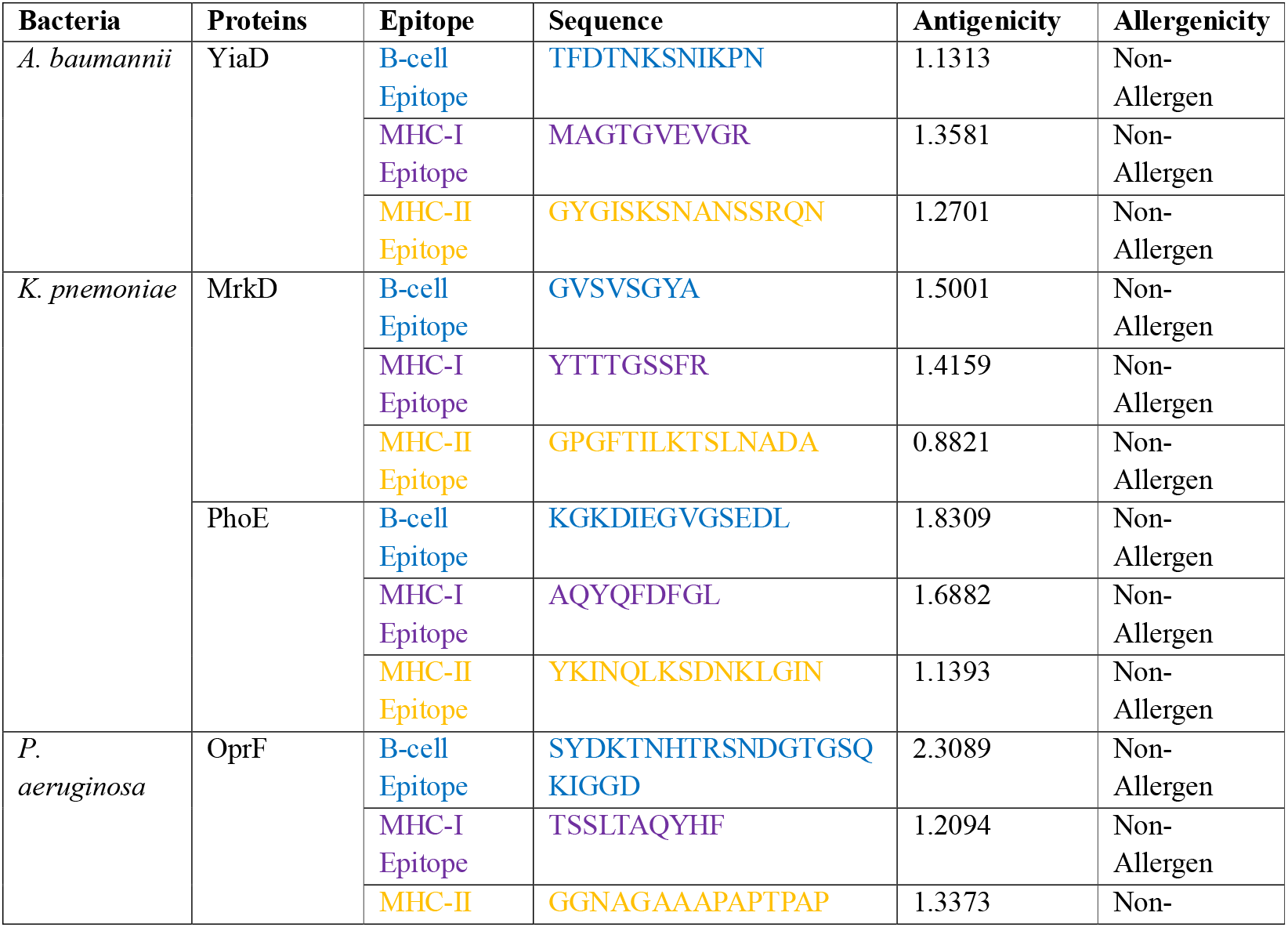
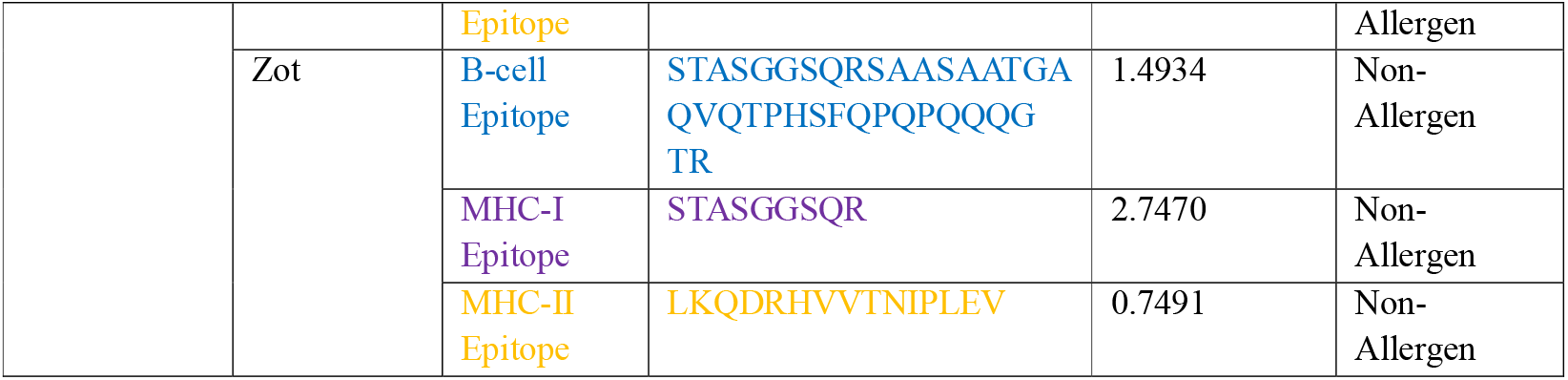
Selected Peptide Epitopes from Immunogenic Proteins.

### Coverage Across Global Population

Population coverage analysis was done for both MHCI and MHCII epitopes combined. In combination, MHC-I and MHC-II epitopes with their alleles demonstrated 98.87% population coverage worldwide. The highest population coverage was found in Europe (99.62%), followed by North America (99.37%), West Indies (98.78%), North East Asia (97.35%), North Africa (96.87%), West Africa (96.64%), Oceania (96.2%), Southeast Asia (95.7%), South Asia (95.5%), Northeast Asia (95.37%), East Africa (94.75%), Central Africa (90.35%), South Africa (93.75%), India (92.26), South America (92.15%), Southwest Asia (91.84%), Central America (53.80), correspondingly (Fig. 1). As a result, the chosen epitopes were determined to be the most promising for creating a multi-epitope peptide vaccine.

**Fig. 1.**
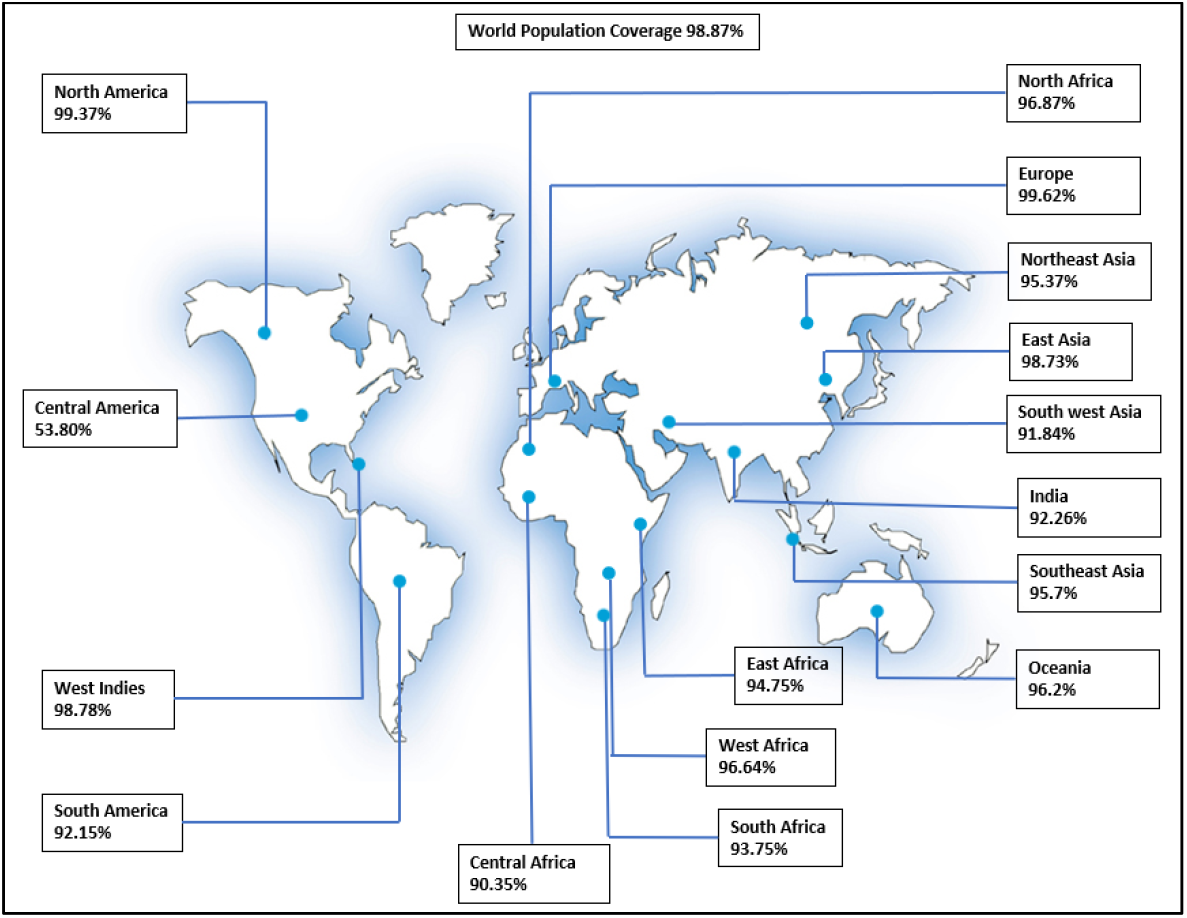
Population coverage of MHC I and MHC II epitopes (combined state)

### Multi-epitope subunit vaccine construction

In the current study, four different adjuvants were incorporated to improve the immunological response. GPGPG was utilized to join five MHC-I epitopes, AAY was used to join five MHC-II epitopes, and KK was used to join B-cell epitopes (Abdus *et al*.,2020). To insert the epitopes with the LptE backbone on one end, EAAAK and other end AAY were added. EAAAK, which gives the sequence a consistent spacing and an alpha helix-forming structure, was used to bind the adjuvant to the MEP construct.

Different B and T-cell epitope sequence patterns were used to obtain the optimal MEP construct, and the final configuration was selected based on antigenicity. Pan DR epitope (PADRE), universal CD4+ cell activator called was combined with 50 ribosomal protein 1.7/L12/ β-defensin/Cholera toxin B subunit/GABA protein/Cholera Toxin B subunit as an adjuvant utilizing an EAAAK linker.

**Fig. 2.**
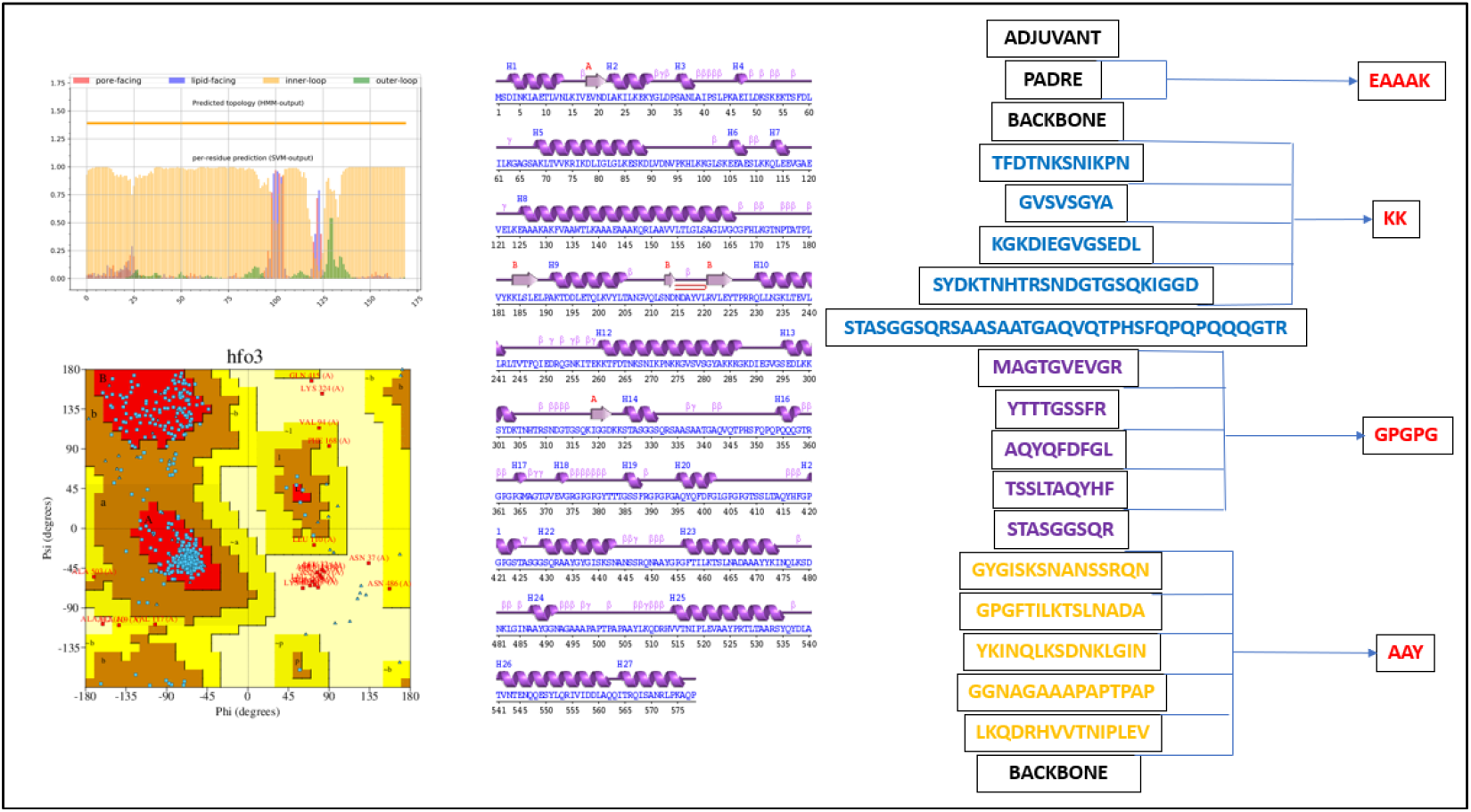
Final MEP construct: a) Topology of LptE b) Ramachandran Plot of MEP c) Secondary structure of MEP d) Final arrangement of MEP construct

### Physicochemical Characterization of MEP

A comparative analysis of four different MEP constructs with their respective adjuvants was done along with the MEP without any adjuvant and based on physicochemical analysis final adjuvant was selected. The best construct for additional analysis among the four adjuvants was 50s ribosomal protein L7/L12, which has the following characteristics: predicted pI value (9.58), aliphatic index (80.95≥61) that confirms stable structure, high antigenicity (0.9253>0.4), high adhesion probability (0.906>0.5), and good water solubility (0.896>0.5) (Supplementary Table S1).

### Computational Modeling and Validation of the Designed Multi-Epitope Subunit Vaccine

SOPMA was used to examine the MEP’s secondary structure (Bibi *et al*., 2021). Final MEP consists of 38.24% alpha helices, 43.60% coils, and 18.17% extended strands. I-TASSER tool was employed to derive the multi-epitope peptide (MEP) construct’s three-dimensional structure (Fig. 3) (Mortazavi et al. 2020). To improve the projected model’s efficiency, the Galaxy Refine web tool was utilized (Abdi et al. 2020). GalaxyRefine generated five different models and enhanced the structure (Supplementary Table S2).

**Fig. 3.**
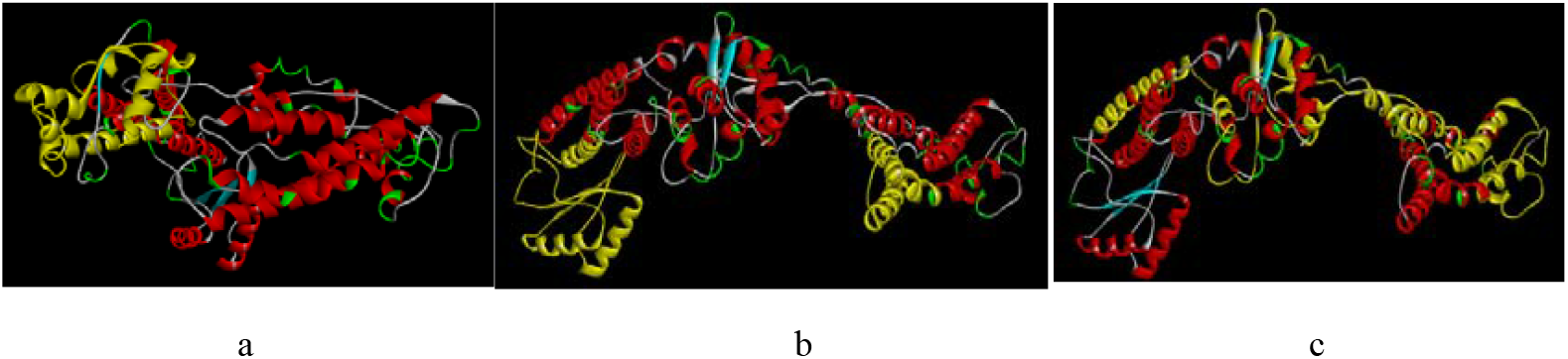
Tertiary Structure of final MEP: a) Adjuvant 50s ribosomal protein L7/L12 (Yellow); b) LptE protein on both ends (Yellow); c) B & T-cell (MHC-I and MHC-II) epitopes (Yellow)

Model 1 was selected for additional processing out of the five anticipated models. The residue percentage in the most favoured area of the Ramachandran plot was 89.4%, the Root Mean Square Deviation (RMSD), MolProbity, and clash score came out to be 0.308 Å, 2.185, and 12.6, respectively, in MEP model 1. The model also displayed less subpar rotamers. The tertiary structure of vaccine model 1 was analysed using a Ramachandran plot with PROCHECK (Rahman et al. 2024). The disallowed region had 13 (2.7%) residues, the generously allowed portions had 1 (1.0%) residue, the additionally allowed regions had 34 (7.0%) residues, and the most favored sections had 436 (89.3%) residues (Table 5). Therefore, these findings confirm the stability of the tertiary structure of MEP, with 97.3% of total residues in permitted regions of the Ramachandran plot.

**Table 5:** Ramachandran Plot Analysis.

| Category | Number of Residues | Percentage (%) |
| --- | --- | --- |
| Most favored regions [A, B, L] | 436 | 89.4 |
| Additional allowed regions [a, b, l, p] | 34 | 7.0 |
| Generously allowed regions [~a, ~b, ~l, ~p] | 5 | 1.0 |
| Disallowed regions [XX] | 13 | 2.7 |
| Total non-glycine, non-proline residues | 488 | 100.0 |
| End residues (excluding Gly and Pro) | 1 | - |
| Glycine residues | 58 | - |
| Proline residues | 31 | - |
| Total residues | 578 | - |

### Prediction of conformational B-cell epitope

Conformational B-cell epitopes if accurately predicted *in silico*, would significantly advance vaccine research, medication formulation, and illness diagnosis (Gabriel et al., 2023). The linear and conformational patterns were evaluated using the Ellipro web server in its default configuration. Ellipro predicted four conformational (discontinuous) and five linear (continuous) B-cell epitopes having antigenicity values ranging from 0.5201 to 2.3063 in the MEP (Supplementary Table S3 and S4, respectively) along with the 3D structure of linear epitopes (Supplementary Fig. 1).

### Disulfide Engineering

For protein engineering, the refined protein model is the best option. Covalent disulfide bonds, which conform to specific geometric constraints, can enhance the structural stability of the model.. Unique disulfide engineering technique can be used to create disulfide bonds with target proteins (Dey et al. 2023). A tool, Disulfide by Design 2.12, was utilised for disulfide bonds generation (Sah et al., 2024). Following server upload, residue pair searching was conducted using the Galaxy-improved phospho-diester bond representation of MEP three-dimensional structure (Gupta et al. 2020). Fifteen latent amino acid pairs were chosen for possible mutation and disulfide engineering out of the forty that were produced (Supplementary Table S5), where cysteine-containing amino acids were the ultimate focus. It has improved the stability of MEP by adding new disulfide links at specific amino acid pairs without changing the MEP structure (Supplementary Figure 1). The mutant construct’s antigenicity was 0.9129, whereas the original construct was 0.9253. With a minor decrease in antigenicity, disulfide engineering improves the MEP construct’s 3-D structure by enhancing its stability (Allemailem., et al. 2021)

### Topology prediction of MEP

The BOCTOPUS program was used to analyze the topology of the final construct. It was found that the outer loop sections of the sequence, which include the majority of the chosen B-cell and T-cell epitopes, are situated between the 250th and 450th amino acids (Supplementary Fig. S3). This confirms the presence of epitopes in MEP construct’s outer loop region.

### Molecular Docking of MEP Construct

MEP and the Toll-like receptors (TLR2 and TLR4) were docked in this study. The Cluspro 2.0 server generated 29 docked complex models (Desta *et al*. 2023) produced 29 models of docked complexes (Supplementary Table S6 & S7). The top 5 models for both the docked complexes are represented in Fig. 4 & 5. The model from TLR2-MEP complex with cluster size 21 and lowest binding energy -1009.6 kcal/mol was selected. Similarly, cluster size and binding energy of 28 and -1079.0 kcal/mol, respectively, were recorded for the best model of the TLR4-MEP docked complex, which was also selected.

**Fig. 4.**
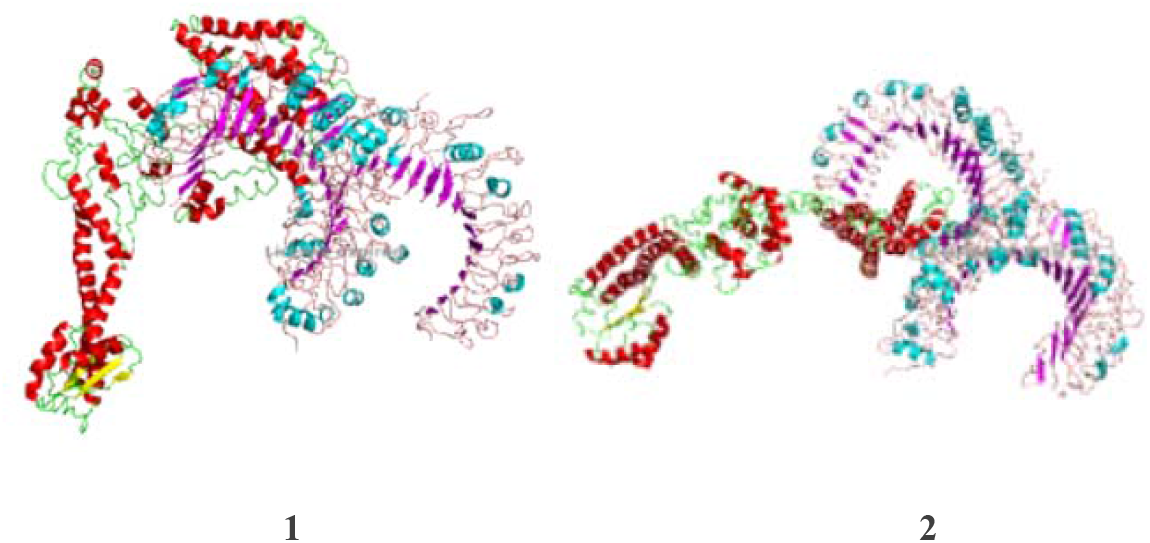
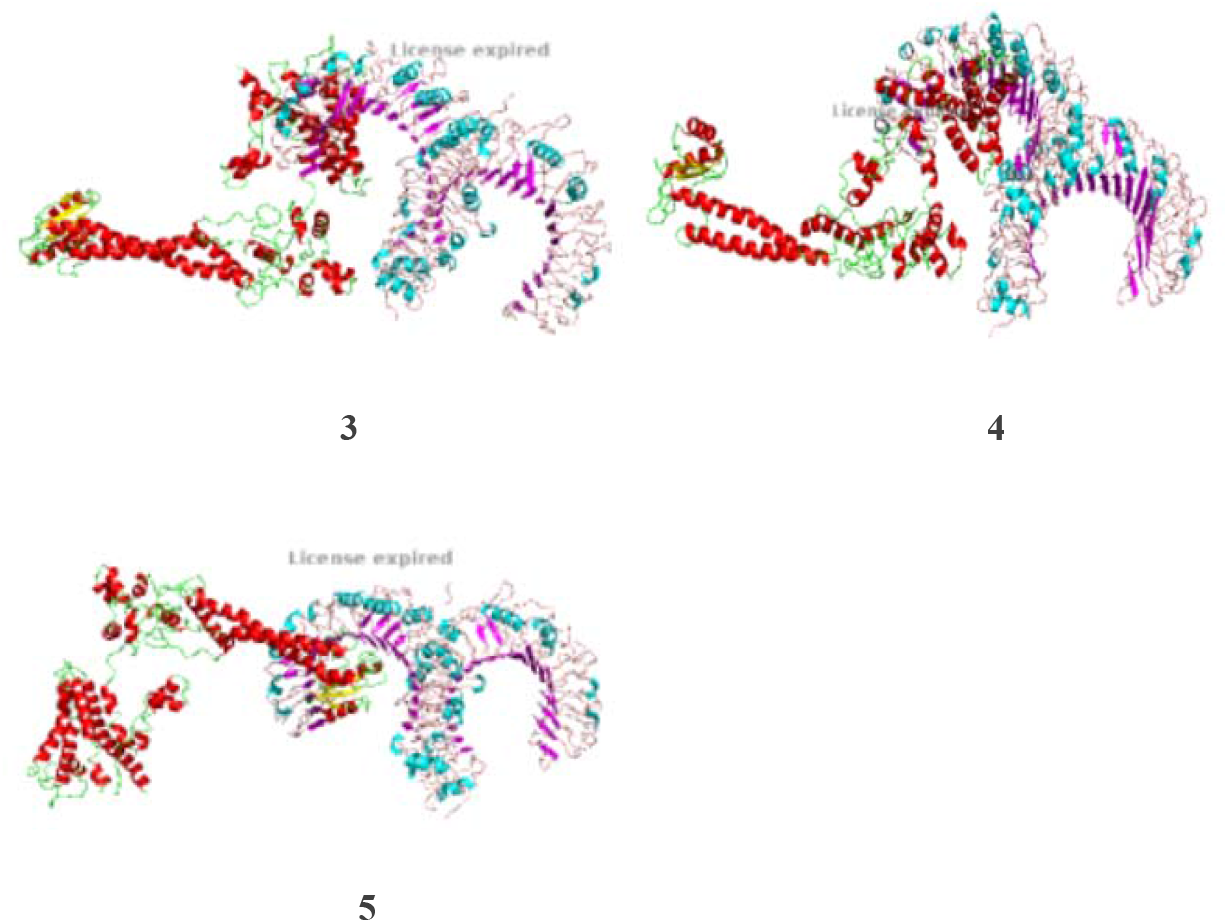
Top 5 models for TLR2-MEP docked complex.

**Fig. 5.**
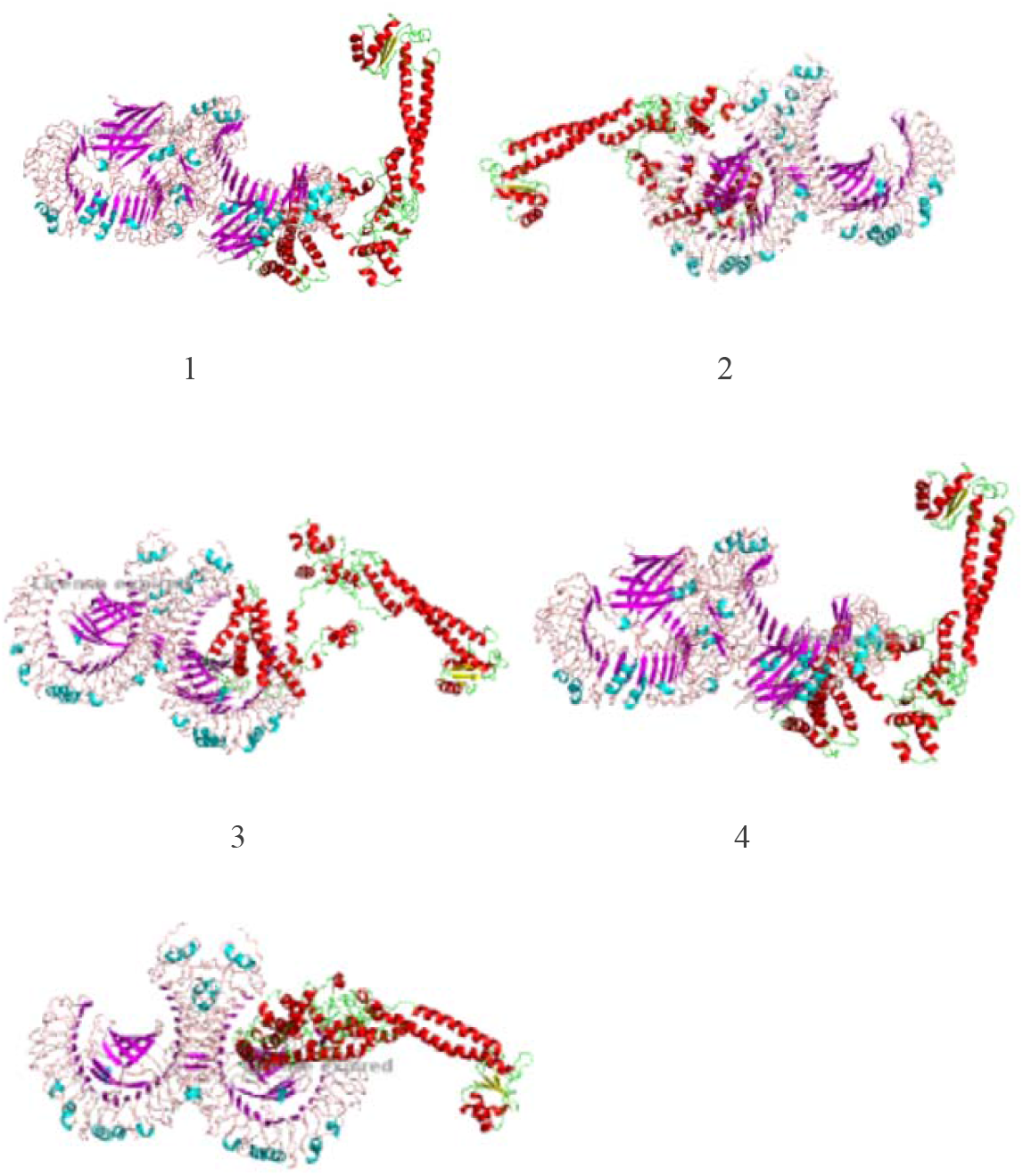
Top 5 models for TLR4-MEP docked complex.

The best models for both MEP docked with TLR-2 and TLR-4 had the highest cluster size and the lowest binding energy.

### Computational Molecular Motion Simulation

Using the iMODS server for molecular dynamics modeling, the stability of the MEP vaccine design was assessed, revealing an improved docked complex with higher molecular flexibility and stability. The MEP-TLR2 and MEP-TLR4 Complexes’ 3D models, Eigenvalues, B-factor/mobility plots, covariance, and variance maps were among the metrics used to achieve this.

**Fig. 3.**
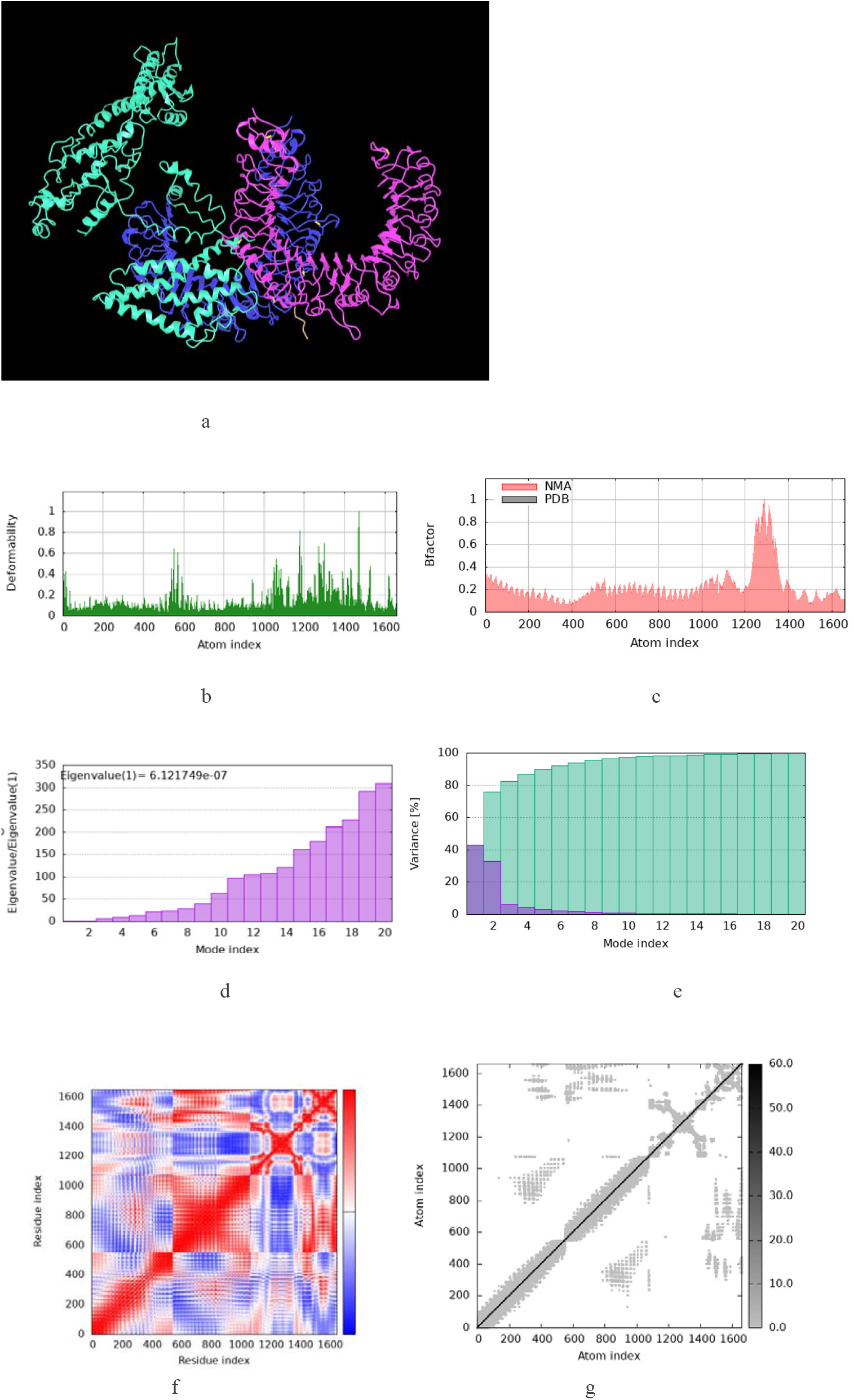
Structural and dynamic analysis of the TLR2–MEP complex: a) Docked conformation of the TLR2–multi-epitope peptide (MEP) complex; b) Deformability profile of the complex; c) B-factor (mobility) plot; d) Eigenvalue graph indicating the complex’s stiffness; e) Variance plot showing individual variance (purple) and cumulative variance (green); f) Covariance matrix plot depicting correlated motions with red, uncorrelated motions with white, and anti-correlated motions with blue; g) Elastic network model representing higher rigidity with darker grey regions

**Fig. 4.**
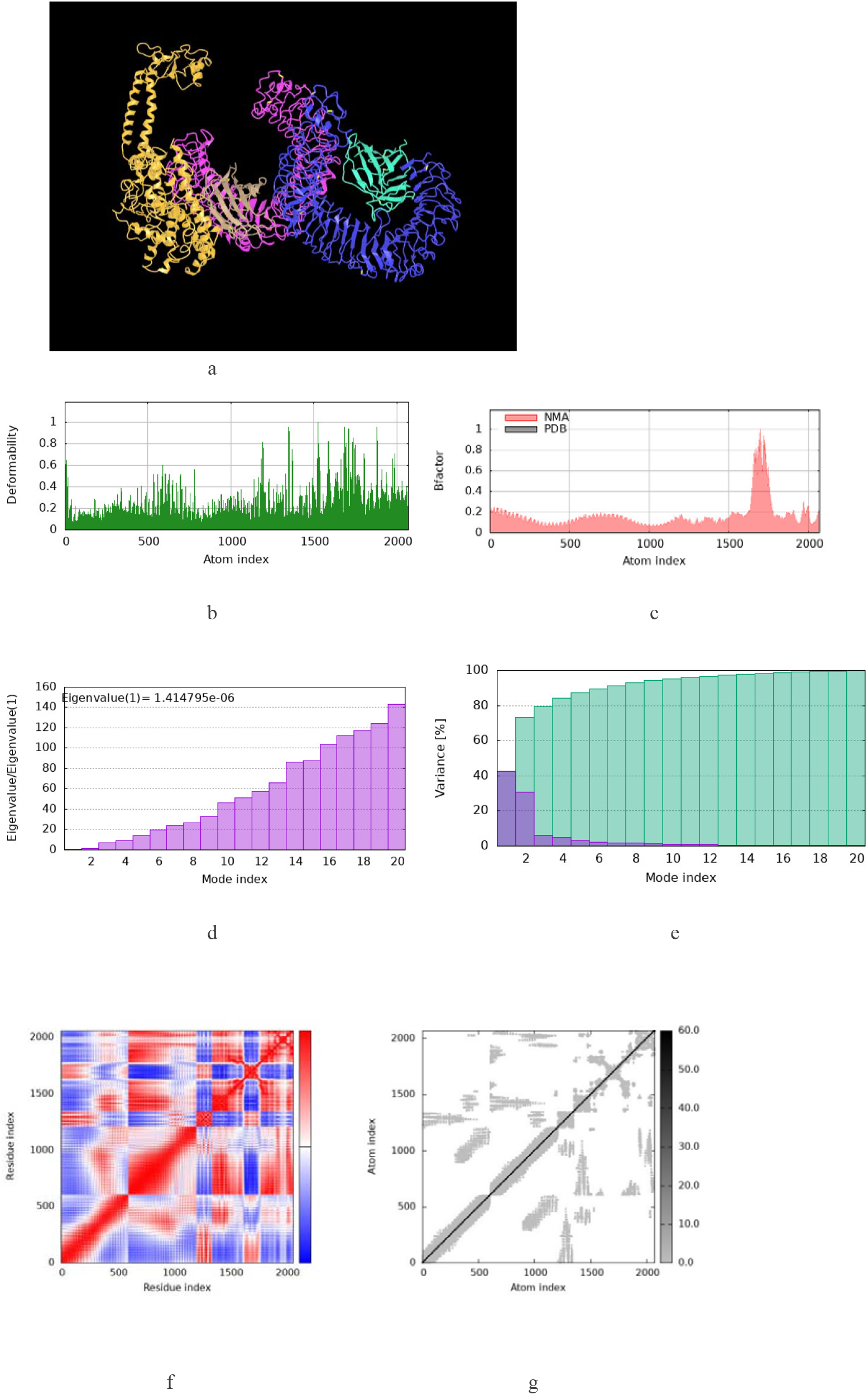
Structural and dynamic analysis of the TLR4–MEP complex: a) Docked structure of the TLR4–multi-epitope peptide (MEP) complex; b) Deformability plot indicating flexible regions; c) B-factor plot representing atomic fluctuations; d) Eigenvalue plot reflecting the stiffness of the complex; e) Variance plot showing individual variance with purple and cumulative variance with green; f) Covariance map illustrating correlated motions with red, uncorrelated motions with white, and anti-correlated motions with blue; g) Elastic network model representing higher rigidity with darker grey regions.

The main-chain deformability graphs for both complexes are used to explore the motions of protein molecules by emphasizing peaks that denote high deformability locations, particularly those around hinge sites.

The variability of each atom allows the B-factor to quantify the MEP vaccine candidate’s adaptability. The B-factor graph shows the relationship between the PDB sector and normal modes analysis of the docked complex. Residue pair correlations are shown in the covariance matrix as follows: blue represents anti-correlated motion, white indicates non-correlated motion, and correlated motion is indicated by red. The RMS values and the B-factor values for normal modes analysis were comparable. To detect atomic pairs connected with springs, elastic network models were developed for the MEP complex with TLR2 and TLR4. The color of each dot in these figures denotes the stiffness of the springs between atomic pairs; stiffer springs are represented by darker gray dots. Through the dynamics of atoms of the MEP docked complex, the ENM reduces the intricate motions of macromolecules to a level that can be understood.

### Codon optimization and In-silico cloning

JCat server and EMBOSS Transeq tool were used for optimizing the codons and reverse translating the sequence in K12 strain of *E. coli*. The translated MEP gene has GC content of 58.93% and CAI value of 0.61. The MEP gene (1734 kb) was inserted into pET28a (+) vector between the restriction sites of BamHI and XhoI for *in-silico* cloning and expression in *E. coli* using SnapGene tool (Supplementary Figure 5). Reverse translation followed by codon optimization were carried out while taking the host’s expression system into account to create the MEP vaccine. GC concentration and codon adaptation index (CAI) were recorded as 51.44%, and 1, respectively, signifying that the gene expression reflected in a GC content ranging from 30% to 70%, as well as a high CAI score (Bibi et al. 2021).

**Fig. 5.**
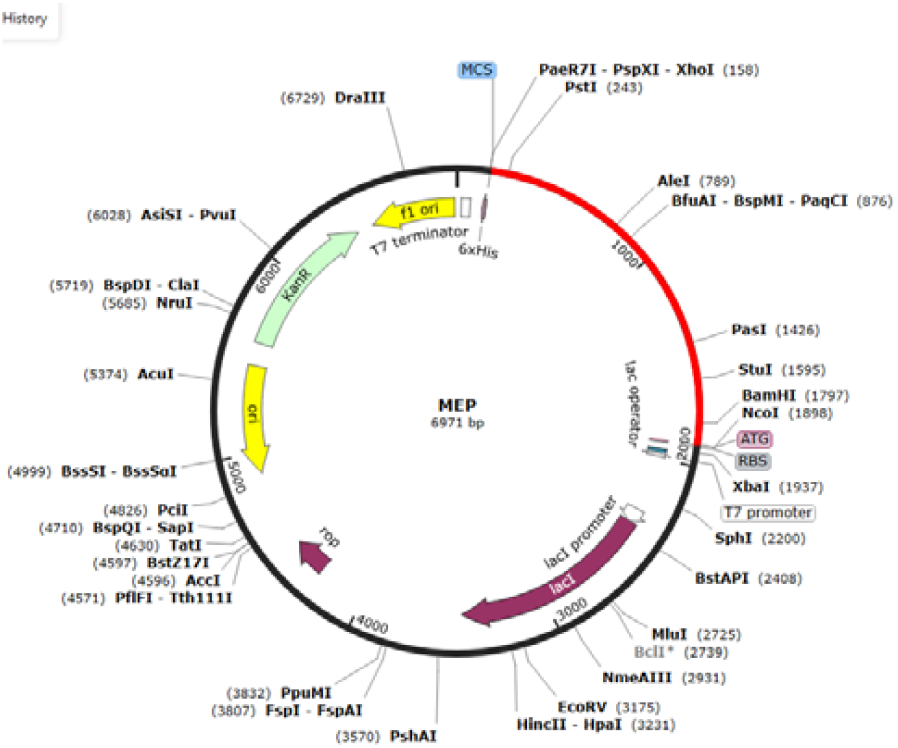
Cloned MEP gene into pET28a.

### C-ImmSim-Based Immune Simulation

Both T-cell and B-cell populations were shown to have significantly increased in *in silico* immunological simulations. Key immunological indicators such asTGF-β, IgM, IgG1, IgG2, IFN-γ, IL-10, and IL-12 were produced as a result of the design of the multiepitope vaccine (MEP), which successfully triggered an immune response. IgM+IgG antibodies were the most prevalent among them, then IgM by itself, followed by IgG1+IgG2, IgG1, and IgG2. The analysis also predicted the expression of various interleukins and cytokines, further supporting the strong immunogenicity and antigenic potential of the MEP construct.

**Fig. 5.**
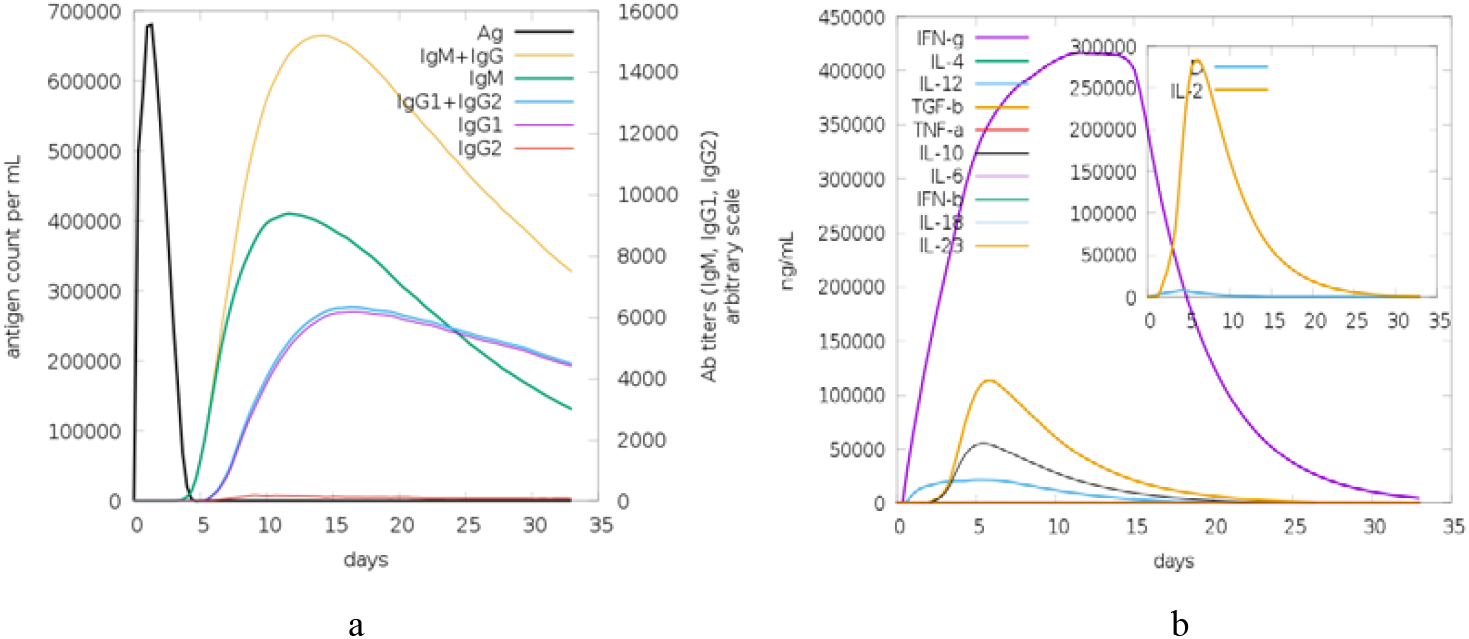
Immune simulation analysis of the MEP construct: a) Graphical representation of antibody production over time in response to antigen exposure; b) Kinetic profile of interleukins and cytokines response

## Discussion

The relentless rise of MDR Gram-negative pathogens, particularly *Acinetobacter baumannii, Klebsiella pneumoniae*, and *Pseudomonas aeruginosa*, poses an unresolved challenge to global healthcare systems. These pathogens, classified as critical or high-priority threats by the WHO, exhibit extensive resistance mechanisms, including β-lactamase production, biofilm formation, and efflux pump activity, rendering conventional antibiotics increasingly ineffective (Pulingam et al., 2022). To get around the drawbacks of conventional vaccine production, this study offers a unique synthetic vaccine based on broad-spectrum epitopes that targets these pathogens. It does this by employing sophisticated immunoinformatics and structural vaccinology techniques.

Traditional vaccine strategies, such as whole-cell inactivated or live-attenuated vaccines, often face challenges related to safety, scalability, and strain-specific immunity (Sharma et al., 2016). Conversely, epitope-based vaccines specifically target immunodominant regions, minimize non-specific response while maximize target specific immune responses (Barman et al., 2022). This approach focused on conserved virulence factors—LptE, YiaD, MrkD, PhoE, OprF, and Zot—that are essential for bacterial survival, and are important virulent factors like adhesion, and immune evasion. By prioritizing these antigens, broad-spectrum coverage was ensured across multiple strains of these pathogens.

The IEDB server was employed to find B and T-cell epitopes that were non-allergenic, highly antigenic, and non-toxic. MHC-I and MHC-II alleles of the chosen epitopes demonstrated strong binding affinities, ensuring robust CD8+ and CD4+ T-cell activation. Notably, the MHC-I epitope from Zot (STASGGSQR) exhibited an exceptionally high antigenicity score (2.7470), suggesting its potential as a dominant immunogen. Population coverage analysis further validated the designed epitope selection, achieving 98.87% global coverage and ensuring effectiveness across diverse populations, including high-coverage regions such as Europe (99.62%) and North America (99.37%). This broad coverage is crucial for a vaccine targeting nosocomial infections, which affect heterogeneous patient populations.

The integration of LptE as a structural backbone provided a scaffold for epitope insertion while maintaining protein stability. The use of flexible linkers (AAY, GPGPG, KK) ensured proper spacing and conformational freedom, critical for epitope presentation (Chen *et al*., 2017). Among the four adjuvant-tested constructs, the 50S ribosomal protein L7/L12 vaccine emerged as the optimal candidate due to its favourable physicochemical properties—high antigenicity (0.9253), solubility (0.665), and structural stability (aliphatic index: 80.95). The refined 3D model exhibited 97.3% residues in the total allowed Ramachandran region, confirming structural integrity.

Docking studies with TLR2 and TLR4 revealed strong binding affinities (TLR2: -1009.6 kcal/mol; TLR4: - 1079.6 kcal/mol), suggesting efficient immune recognition. The large cluster sizes (21 for TLR2, 28 for TLR4) indicated stable interactions, a prerequisite for effective vaccine-induced immunity. Molecular dynamics simulations confirmed these findings, demonstrating low deformability and high rigidity in key binding regions, indicative of a stable vaccine-receptor complex.

*In silico* immune simulations predicted robust humoral and cellular responses, with elevated IgM/IgG levels and Th1/Th2 cytokine profiles (IFN-γ, IL-12). The rapid IgM surge post-exposure, followed by sustained IgG1/IgG2 production, depicts natural immune responses to bacterial infections, suggesting that the vaccine could provide both immediate as well as long-lasting protection. The IFN-γ and IL-12 induction further supports a Th1-polarized response, essential for combating intracellular pathogens.

Currently, there is no vaccine for *A. baumannii/K. pneumoniae/P. aeruginosa* exists, and a single broad-spectrum synthetic MEP vaccine against these three pathogens is a novel approach towards next-generation vaccine development by targeting multiple conserved antigens simultaneously. Additionally, the inclusion of a PADRE sequence enhances CD4+ T-cell activation, addressing a common limitation of peptide-based vaccines.

The *in-silico* results from this study demonstrate strong potential, while further *in vitro* and *in vivo* validations are essential. Challenges such as epitope processing efficiency, potential immune evasion, and adjuvant compatibility must be addressed. However, this MEP design can be further improved by incorporating additional adjuvants or epitopes based on experimental feedback. If successful, this vaccine could revolutionize prophylactic strategies in high-risk settings (ICUs, burn units), significantly reducing MDR infection rates.

## Conclusion

The increasing threat of multidrug-resistant (MDR) Gram-negative bacteria, particularly *Acinetobacter baumannii, Klebsiella pneumoniae*, and *Pseudomonas aeruginosa*, necessitates new and proactive approaches beyond conventional antibiotics. This work presents a new broad-spectrum, multi-epitope synthetic vaccine developed by cutting-edge immunoinformatics and structural vaccinology approaches. Targeting strongly conserved and immunogenic epitopes of key bacterial proteins—LptE, YiaD, MrkD, PhoE, OprF, and Zot—a vaccine construct that can elicit robust and persistent immune responses against these notorious pathogens was developed. By incorporating computational predictions with experimental confirmations, the way for next-generation vaccines that are specific, broadly protective, and responsive to emerging bacterial threats was paved.

