## Supplementary file for "Immunoinformatic Designing and Evaluation of a Broad-Spectrum Multiepitope Vaccine Against MDR *Acinetobacter baumannii, Klebsiella pneumoniae* and *Pseudomonas aeruginosa*"

**Supplementary Material:**

**Table S1: Comparative analysis of vaccine constructs without adjuvant and with different adjuvants**

| **Property** | **Without Adjuvant** | **With Adjuvant** | | | |
| --- | --- | --- | --- | --- | --- |
|  |  | **50s Ribosomal protein** | **Beta-defensin** | **GABA protein** | **Cholera toxin** |
| Amino acid | 426 | 578 | 500 | 613 | 578 |
| Mol. Wt. | 44864.04 | 61237.32 | 52970.76 | 65307.02 | 61621.48 |
| Aliphatic Index | 70.16 | 80.95 | 69.98 | 75.77 | 74.57 |
| Antigenicity | 1.0897 | 0.9253 | 1.0453 | 0.9491 | 0.9662 |
| Allergenicity | Non allergen | Non allergen | Non allergen | Non allergen | Non allergen |
| Virulency | Virulent | Virulent | Virulent | Virulent | Virulent |
| Toxicity | Non-Toxin | Non-Toxin | Non-Toxin | Non-Toxin | Non-Toxin |
| Solubility | 0.452 | 0.665 | 0.880 | 0.565 | 0.863 |
| Hydropathicity | -0.774 | -0.479 | -0.546 | -0.526 | -0.458 |
| pI | 9.67 | 9.58 | 9.82 | 9.26 | 9.60 |
| Refined Ramachandran  plot | 85.0 % | 89.3% | 75.5% | 84.2% | 86.6% |

**Table S2: Refined models of MEP**

| **Model** | **GDT-HA** | **RMSD** | **MolProbity** | **Clash score** | **Poor rotamers** | **Rama favored** |
| --- | --- | --- | --- | --- | --- | --- |
| Initial | 1.0000 | 0.000 | 2.300 | 13.5 | 1.5 | 91.0 |
| MODEL 1 | 0.9853 | 0.308 | 2.185 | 12.6 | 1.3 | 92.2 |
| MODEL 2 | 0.9849 | 0.306 | 2.208 | 13.2 | 1.3 | 92.2 |
| MODEL 3 | 0.9840 | 0.315 | 2.054 | 11.7 | 0.4 | 92.4 |
| MODEL 4 | 0.9745 | 0.328 | 2.083 | 12.6 | 0.4 | 92.4 |
| MODEL 5 | 0.9844 | 0.310 | 2.142 | 14.1 | 0.9 | 92.0 |

**Table S3**: Discontinuous B-cell Epitopes

| No. | Residues | No. of Residues | Score | Antigenicity |
| --- | --- | --- | --- | --- |
| 1 | A:L157,A:G158,A:L159,A:S160,A:A161,A:G162,A:L163,A:V164,A:G165, A:C166, A:G167, A:F168, A:H169, A:L170, A:K171, A:G172, A:T173, A:N174, A:P175, A:T176, A:A177, A:T178, A:P179, A:L180, A:V181, A:Y182, A:K183, A:K184, A:L185, A:S186, A:L187, A:E188, A:L189, A:P190, A:A191, A:K192, A:T193, A:D194, A:D195, A:L196, A:E197, A:T198, A:Q199, A:L200, A:K201, A:V202, A:Y203, A:L204, A:T205, A:A206, A:N207, A:G208, A:V209, A:Q210, A:L211, A:S212, A:N213, A:D214, A:N215, A:D216, A:A217, A:Y218, A:V219, A:L220, A:R221, A:V222, A:L223, A:E224, A:Y225, A:T226, A:P227, A:R228, A:R229, A:Q230, A:L231, A:L232, A:N233, A:G234, A:K235, A:L236, A:T237, A:E238, A:V239, A:L240, A:L241, A:R242, A:L243, A:T244, A:V245, A:T246, A:F247, A:Q248, A:I249, A:E250, A:D251, A:R252, A:G254, A:N255, A:K256, A:I257, A:T258, A:E259, A:K260, A:K261, A:T262, A:F263, A:D264, A:T265, A:N266, A:K267, A:S268, A:N269, A:I270, A:N273 | 114 | 0.784 | **0.5201** |
| 2 | A:T369, A:T409, A:S410, A:S411, A:L412, A:T413, A:A414, A:Q415, A:Y416, A:H417, A:F418, A:G419, A:P420, A:G421, A:P422, A:G423, A:S424, A:T425, A:A426, A:Y453, A:L460, A:K461, A:S463, A:L464, A:N465, A:A466, A:D467, A:A468, A:A469, A:A470, A:Y471, A:Y472, A:K473, A:I474, A:N475, A:Q476, A:L477, A:K478, A:S479, A:D480, A:N481, A:K482, A:L483, A:G484, A:I485, A:N486, A:A487, A:A488, A:Y489, A:G490, A:G491, A:N492, A:A493, A:G494, A:A495, A:A496, A:A497, A:P498, A:A499, A:P500, A:T501, A:P502, A:A503, A:P504, A:A505, A:A506, A:Y507, A:L508, A:K509, A:Q510, A:D511, A:R512, A:H513, A:V514, A:V515, A:T516, A:N517, A:I518, A:P519, A:L520, A:E521, A:V522, A:A523, A:A524, A:Y525, A:P526, A:R527, A:Q563, A:R566, A:Q567, A:A570, A:N571, A:L573, A:P574, A:K575, A:A576, A:Q577, A:P578 | 98 | 0.728 | **0.5963** |
| 3 | A:I16, A:V17, A:E18, A:V19, A:N20, A:D21, A:L22, A:A23, A:K24, A:I25, A:L26, A:K27, A:E28, A:K29, A:Y30, A:G31, A:L32, A:D33, A:P34, A:S35, A:A36, A:N37, A:L38, A:A39, A:I40, A:P41, A:S42, A:L43, A:P44, A:K45, A:A46, A:E47, A:I48, A:L49, A:D50, A:K51, A:S52, A:K53, A:E54, A:K55, A:T56, A:S57, A:F58, A:D59, A:L60, A:I61, A:L62, A:K63, A:G64, A:A65, A:G66, A:S67, A:A68, A:K69, A:V117, A:G118, A:A119, A:E120, A:V121, A:E122, A:L123, A:K124, A:N306, A:H307, A:S310, A:N311, A:D312, A:G313, A:T314, A:K318, A:I319, A:G320, A:G321, A:D322, A:K323, A:K324, A:S325, A:T326, A:A327, A:Q345 | 80 | 0.643 | **0.7671** |
| 4 | A:S447, A:R448, A:Q449, A:L529, A:T530, A:A531, A:A532, A:R533, A:S534, A:Y535, A:Q536, A:D538, A:L539, A:A540, A:T541, A:V542, A:E545 | 17 | 0.554 | **0.5784** |

**Table S4**: Continuous B-cell Epitopes

| No. | Start | End | Peptide | No. of Residues | Score | Antigenicity |
| --- | --- | --- | --- | --- | --- | --- |
| 1 | 157 | 270 | LGLSAGLVGCGFHLKGTNPTATPLVYKKLSLELPAKTDDLETQLKVYLTANGV  QLSNDNDAYVLRVLEYTPRRQLLNGKLTEVLLRLTVTFQIEDRQGNKITEKKT  FDTNKSNI | 114 | 0.784 | **0.8545** |
| 2 | 460 | 542 | LKTSLNADAAAYYKINQLKSDNKLGINAAYGGNAGAAAPAPTPAPAAYLKQD  RHVVTNIPLEVAAYPRTLTAARSYQYDLATV | 83 | 0.726 | **0.6911** |
| 3 | 16 | 69 | IVEVNDLAKILKEKYGLDPSANLAIPSLPKAEILDKSKEKTSFDLILKGAGSAK | 54 | 0.724 | **NON-ANTIGEN** |
| 4 | 409 | 426 | TSSLTAQYHFGPGPGSTA | 18 | 0.692 | **1.3781** |
| 5 | 117 | 124 | VGAEVELK | 8 | 0.532 | **2.3063** |

**Table S5:** Amino acid pairs for potential mutation and disulfide engineering with selected amino acids (Green) and excluded amino acids (Red) for disulphide bond formation

|  | **Residue 1** | | | **Residue 2** | | | **Bond** | | |
| --- | --- | --- | --- | --- | --- | --- | --- | --- | --- |
|  | Chain | Position | Amino Acid | Chain | Position | Amino Acid | **χ_3_** | kcal/mol | ƩB-factor |
|  | A | 144 | ALA | A | 278 | SER | -116.64 | 4.04 | 0.00 |
|  | A | 468 | ALA | A | 489 | TYR | -107.04 | 3.25 | 0.00 |
|  | A | 110 | LEU | A | 293 | VAL | -106.97 | 3.00 | 0.00 |
|  | A | 496 | ALA | A | 499 | ALA | -101.40 | 4.15 | 0.00 |
|  | A | 89 | ASP | A | 137 | THR | -98.46 | 1.02 | 0.00 |
|  | A | 449 | GLN | A | 535 | TYR | -92.30 | 1.96 | 0.00 |
|  | A | 170 | LEU | A | 175 | PRO | -90.91 | 5.05 | 0.00 |
|  | A | 492 | ASN | A | 502 | PRO | -89.98 | 3.32 | 0.00 |
|  | A | 475 | ASN | A | 480 | ASP | -79.33 | 451 | 0.00 |
|  | A | 481 | ASN | A | 503 | ALA | -71.92 | 5.88 | 0.00 |
|  | A | 34 | PRO | A | 37 | ASN | -71.82 | 3.92 | 0.00 |
|  | A | 429 | GLY | A | 562 | GLN | -71.33 | 4.60 | 0.00 |
|  | A | 14 | LEU | A | 348 | HIS | -71.13 | 4.52 | 0.00 |
|  | A | 174 | ASN | A | 177 | ALA | -68.96 | 7.03 | 0.00 |
|  | A | 101 | GLY | A | 332 | GLN | -64.69 | 5.72 | 0.00 |
|  | A | 69 | LYS | A | 307 | HIS | +84.60 | 4.61 | 0.00 |
|  | A | 62 | LEU | A | 75 | ARG | +85.31 | 4.16 | 0.00 |
|  | A | 334 | SER | A | 349 | SER | +88.62 | 1.88 | 0.00 |
|  | A | 99 | LYS | A | 107 | ALA | +9096 | 3.86 | 0.00 |
|  | A | 393 | PRO | A | 397 | TYR | +92.80 | 4.63 | 0.00 |
|  | A | 301 | SER | A | 343 | GLN | +92.80 | 0.37 | 0.00 |
|  | A | 378 | GLY | A | 406 | GLY | +93.28 | 4.26 | 0.00 |
|  | A | 449 | GLN | A | 531 | ALA | +94.04 | 5.73 | 0.00 |
|  | A | 14 | LEU | A | 330 | GLY | +94.04 | 3.99 | 0.00 |
|  | A | 479 | SER | A | 505 | ALA | +94.25 | 5.31 | 0.00 |
|  | A | 367 | ALA | A | 568 | ILE | +95.04 | 4.60 | 0.00 |
|  | A | 26 | LEU | A | 51 | LYS | +95.50 | 2.61 | 0.00 |
|  | A | 481 | ASN | A | 485 | ILE | +100.28 | 2.98 | 0.00 |
|  | A | 469 | ALA | A | 486 | ASN | +104.09 | 2.97 | 0.00 |
|  | A | 58 | PHE | A | 68 | ALA | +108.15 | 7.37 | 0.00 |
|  | A | 413 | THR | A | 419 | GLY | +110.30 | 4.82 | 0.00 |
|  | A | 411 | SER | A | 569 | SER | +110.79 | 3.85 | 0.00 |
|  | A | 104 | LYS | A | 107 | ALA | +115.93 | 2.96 | 0.00 |
|  | A | 488 | ALA | A | 503 | ALA | +117.06 | 6.51 | 0.00 |
|  | A | 83 | GLY | A | 292 | GLY | +117.15 | 5.37 | 0.00 |
|  | A | 59 | ASP | A | 305 | THR | +117.99 | 5.80 | 0.00 |
|  | A | 102 | LEU | A | 328 | SER | +119.22 | 3.12 | 0.00 |
|  | A | 21 | ASP | A | 24 | LYS | +123.28 | 3.24 | 0.00 |
|  | A | 447 | SER | A | 536 | GLN | +124.89 | 5.41 | 0.00 |
|  | A | 66 | GLY | A | 71 | THR | +125.79 | 5.17 | 0.00 |

**Table S6: Cluster members and binding energies for top 5 generated models of TLR 2-MEP Docked Complex**

| **Cluster** | **Members** | **Representative** | **Weighted Score** |
| --- | --- | --- | --- |
| **1** | 21 | Center | -832.9 |
|  |  | Lowest Energy | -1009.6 |
| **2** | 16 | Center | -825.9 |
|  |  | Lowest Energy | -987.3 |
| **3** | 23 | Center | -768.7 |
|  |  | Lowest Energy | -938.2 |
| **4** | 30 | Center | -796.1 |
|  |  | Lowest Energy | -925.8 |
| **5** | 39 | Center | -765.8 |
|  |  | Lowest Energy | -908.7 |

**Table S7: Cluster members and binding energies for top 5 generated models of TLR 4-MEP Docked Complex**

| **Cluster** | **Members** | **Representative** | **Weighted Score** |
| --- | --- | --- | --- |
| **1** | 21 | Center | -963.1 |
|  |  | Lowest Energy | -1089.0 |
| **2** | 28 | Center | -812.3 |
|  |  | Lowest Energy | -1079.1 |
|  |  | Lowest Energy | -935.8 |
| **3** | 22 | Center | -984.5 |
|  |  | Lowest Energy | -1064.0 |
| **4** | 21 | Center | -1007.8 |
|  |  | Lowest Energy | -1061.8 |
|  |  | Lowest Energy | -991.9 |
| **5** | 18 | Center | -820.8 |
|  |  | Lowest Energy | -1035.3 |

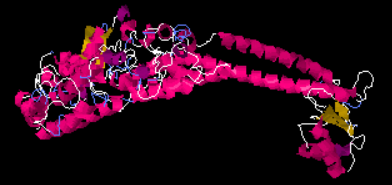

Original

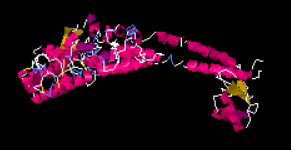

Mutant

**Fig. S1** 3D structure of Mep before disulfide engineering (Original) and after disulfide engineering (Mutant)

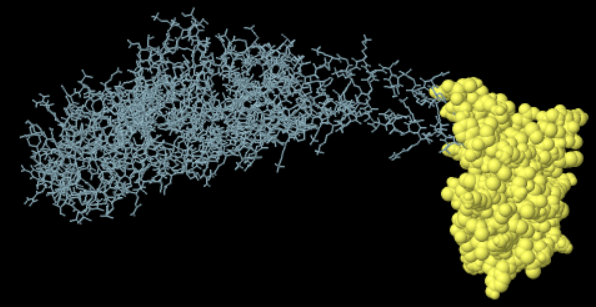

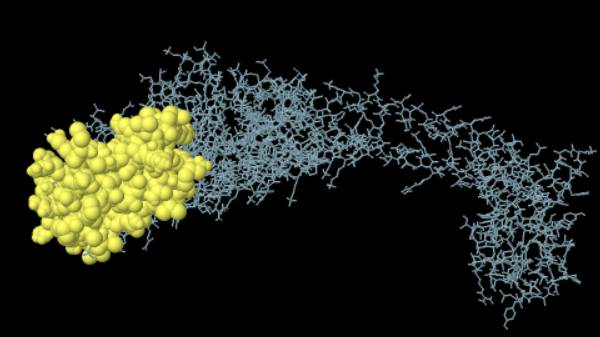

1	2

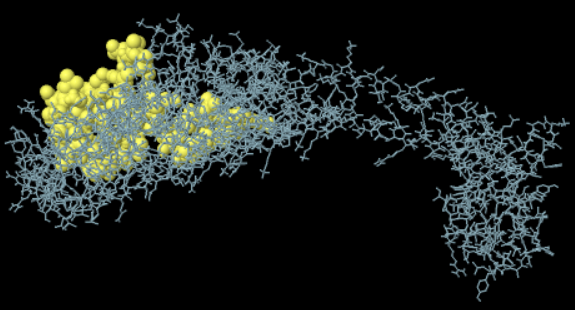

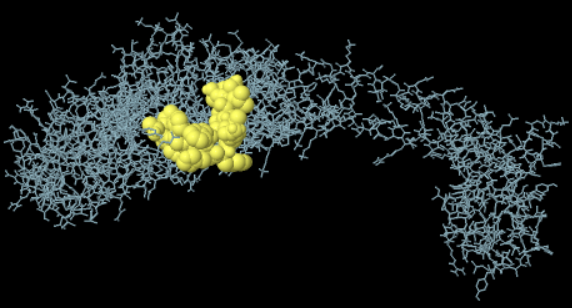

3	4

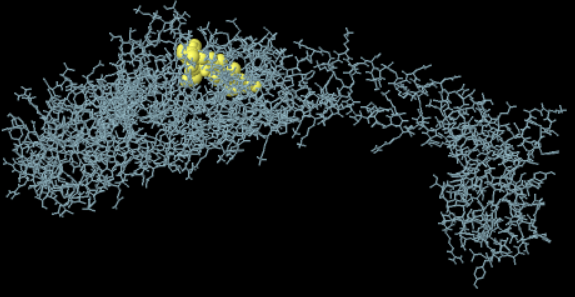

5

**Fig**. **S2** Continuous B-cell Epitopes: 3D structure of 1,2,3,4 and 5 residues of continuous epitopes

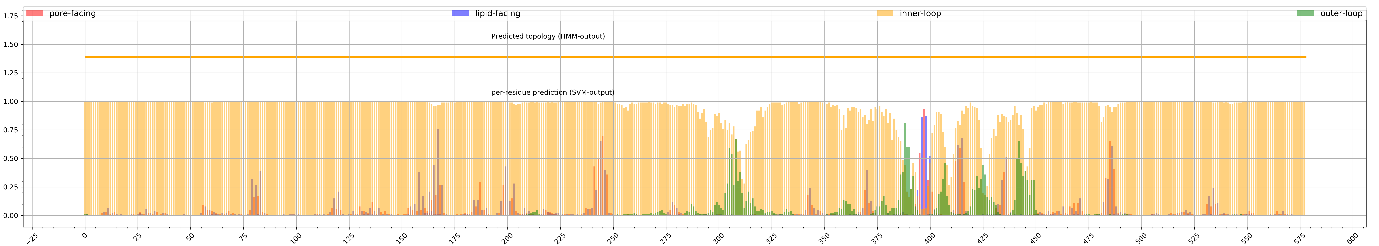

**Fig. S3** Topology of final MEP construct
